# Novel two-stage deep learning-based approach applied to gene expression data pertaining to esophageal adenocarcinoma boosting biological knowledge discovery

**DOI:** 10.64898/2026.09.12.751181

**Authors:** Fahad Jamie, Turki Turki, Fawaz Alsolami, Y-h. Taguchi

## Abstract

Esophageal cancer (EC) is characterized by complex transcriptional alterations and therapeutic resistance, posing challenges for traditional computational methods. In this study, we propose a deep learning (DL)-based computational framework to identify important genes and biologically relevant pathways in bulk cell RNA-seq data (GSE234304 and GSE273848), which comprise tumor and non-tumor esophageal tissue samples. A fully connected feedforward neural network was trained for binary classification, and two feature selection strategies were implemented: Neural Network followed by Support Vector Regression (NN+SVR) and Integrated Gradients combined with SVR (IG+SVR). The genes were then ranked according to their weights in SVR deriving the importance scores, and the top 100 genes were subjected to enrichment analysis using Enrichr and Metascape. The proposed DL-based approaches identified a greater number of expressed genes across established esophageal cancer cell lines than LIMMA, SAM, and the t-test did. Specifically, in the GSE234304 dataset, IG + SVR, our best method, identified a total of 9 expressed genes while the best baseline method, LIMMA, identified a total of 3 expressed genes. In terms of GSE273848 dataset, IG + SVR was also the best identifying a total of 11 expressed genes while the best baseline method, t-test, had a total of 7 expressed genes. The key genes identified included CEBPB, SUMO1, RORA, STAT1, GATA, OCT1, RUNX1, and NR3C1, as well as pathways related to nucleoprotein maturation, collagen fibril organization, the immunoglobulin-mediated immune response, immune regulation, and insulin signaling. These results show that combining neural networks and attribution-based regression creates an effective and interpretable framework for selecting genes in esophageal cancer research.

## 1. Introduction

Esophageal cancer (EC) is associated with poor prognosis and can cause deaths if metastasized and not detected at earlier stages [1]. Risk factors including hypertension, smoking, obesity, and lifestyle, to just name a few, can contribute to increasing the likelihood of getting EC [2], [3], [4]. Researchers have developed computational methods to uncover various molecular mechanisms underlying EC. Hasib et al. [5] identified hub genes and lncRNA pertaining to esophageal squamous cell carcinoma (ESCC), following baseline-based approach. First, they downloaded three gene expression data (GSE161533, GSE20347, and GSE45670) from the gene expression omnibus (GEO). They employed the GEO2R tool to the gene expression data of normal and ESCC samples, identifying 75 DEGs that were then provided as input to DAVID to perform enrichment analysis. STRING database was used to identify the protein-protein interaction network, identifying 10 hub genes (PLAU, CXCL1, KIF14, MMP1, SPP1, CXCL8, MMP3, MMP13, UBE2C, and TOP2A). Kaplan-Meier analysis was used to identify five hub genes (MMP13, CXCL8, TOP2A, CXCL1 and SPP1). HYMAI was identified by HUGO database as a expressed among the three datasets. Zhang et al. [6] integrated single-cell RNA-seq and bulk transcriptomic data to classify ESCC tumors into four molecular subtypes and identify subtype-specific marker genes. Using Cox and LASSO regression, they constructed a four-gene prognostic signature (CCND1, PKP1, JUP, ANKRD12) capable of predicting overall survival with an AUC of approximately 0.73.

Xie et al. [7] identified important cancer-promoting genes pertaining to drug resistance in ESCC through the use of a bioinformatics-based approach. First, they downloaded GSE86099 and GSE50224 from the GEO, in addition to TCGA-ESCA. They applied DEGseq, identifying 2942 DE mRNAs in GSE86099. Then, identifying 35 common genes as oncogenes (including DOCK8, PTPRD, and RBM23). The PPI network included eight hub genes. Five genes (BMP1, DBF4, HIP1, ANG, MAP7D2) were related to ESCC resistance. Four genes (BMP1, HIP1, ANG, MAP7D2) were used with lasso, unveiling molecular mechanism of ESCC. He et al. [8] analyzed GEO transcriptomic data before and after radiotherapy in EC and identified 28 DEGs associated with treatment response. Using LASSO and SVM, they derived a three-gene signature (ZDHHC11B, CD46, PFKFB3) associated with immune microenvironment changes and increased CD8⁺ T-cell infiltration after radiotherapy. Yuan et al. [9] obtained gene expression data and miRNA-seq related to normal and tumor samples from the TCGA database to identify novel miRNA markers. They identified 1319 DEGs from the gene expression data while identifying target genes related to 48 differentially expressed miRNAs (DEMs). There were four hundred common DEGs, provided to GO and KEGG for enrichment analysis, followed by identifying 66 genes in the biological pathways. Then, miRNA-mRNA network was constructed. Nine DEMs and 28 target genes were identified. Finally, miRNA-mRNA network was constructed from 9 DEMs and 28 targets in the biological pathways, yielding three miRNA markers (hsa-miR-452-3p, hsa-miR-6499-3p,hsa-miR-205-3p) related to the first three stages of ESCC.

Khurshid et al. [10] identified hub genes acting as potential biomarkers and therapeutic targets for ESCC through the use of a bioinformatics-based approach. Data were downloaded from TCGA and GTEx databases. Limma was applied reporting 2524 DEGs between TCGA-ESCC samples and GTEx normal samples. A pathway enrichment analysis resulted in 365 genes, provided as input to STRING to construct the PPI network. Twenty hub genes were reported in which CDK1 MAD2L1, PLK1, and TOP2A were among those genes. Fu et al. [11] identified PANoptosis-related genes (PRGs), acting as a gene signature for Esophageal adenocarcinoma (EAC). Eighty-eight EAC samples were downloaded from TCGA-ESCA while the other 652 samples pertaining to normal esophageal were downloaded from GTEx. Limma was applied to the samples, identifying 8776 DEGs. There was 273 DEGs resulted from the intersection of 8776 DEGs with 624 PRGs from previous studies. Univariate Cox regression was then utilized to gene expression with 624 genes, yielding a model with 18 genes. Lasso was then utilized to identify eight genes including ATRX, ERBB2, MSLN, MMP12, CLU, COL11A1, TERT, PSMA1. The eight genes was validated using GSE13898 data from the GEO database, reaching an AUC values of 0.766, 0.682, and 0.694 for 5-year survival rate, 3-year survival rate, and 1-year survival rate, respectively. Weighted Gene Co-expression Network Analysis (WGCNA) identified these hub genes: i MMP12, RBB2, and CLU.

Zhang et al. [12] investigated cuproptosis-related genes in ESCC and used LASSO-Cox regression to construct an 11-gene prognostic signature based on 12 candidate genes. Patients classified in the high-risk group exhibited significantly poorer overall survival compared with the low-risk group. Shi et al. [13] integrated single-cell RNA-seq and bulk transcriptomic data to identify tumor stemness characteristics in esophageal cancer and constructed an 18-gene tumor stem cell marker signature using LASSO-Cox regression. The resulting risk model revealed significant differences in immune infiltration and chemotherapy sensitivity between risk groups.

Alghamdi et al. [14] proposed a deep learning–based framework for analyzing single-cell RNA-seq data to identify genes associated with drug resistance in melanoma. Using two GEO scRNA-seq datasets (GSE108383_A375 and GSE108383_451Lu), a fully connected neural network combined with explainability methods (DeepLIFT, SHAP, Integrated Gradients, and LRP) identified 100 important genes, including ARAF, SOX10, DCT, and AXL, and detected melanoma-related drugs such as Vemurafenib and Dabrafenib, while Integrated Gradients improved classification performance by 2.2% and 0.5% over baseline methods. Zhu et al. [15] constructed a machine learning-based prognostic model for ESCC based on 15 glycolipid metabolism-related genes using TCGA and GEO datasets. The model stratified patients into high- and low-risk groups with significantly different survival outcomes and revealed differences in immune cell infiltration patterns between the two groups. Zhao et al. [16] employed RNA-seq and machine learning to identify genes associated with lymph node metastasis (LNM) in ESCC. RNA-seq analysis identified 2,837 DEGs between LNM-positive and LNM-negative tumors, and a random forest model highlighted SIM2, CUX1, and CYP4B1 as predictive markers, while a logistic regression model integrating these genes with clinical features achieved an AUC of 0.83 for LNM prediction.

Bioinformatics methods remain central to investigating drug mechanisms in esophageal cancer; however, they predominantly depend on established analytical tools and make limited use of advanced artificial intelligence techniques. In addition, current AI-based approaches have not fully explored the integration of deep learning (DL) with gene expression data analysis and therefore do not yet capture the full potential of these combined methodologies.

In this study, we propose a computational framework integrating adapted deep learning (DL) models with downstream enrichment analysis. This framework is specifically tailored for analyzing gene expression data. This design allows for a more detailed characterization of underlying biological processes. The framework is also intended to facilitate the translation of computational findings into clinical contexts by identifying therapeutic targets associated with favorable treatment responses in esophageal cancer. Unlike previous studies that relied on linear statistical assumptions, our framework addresses the nonlinear transcriptional heterogeneity characteristic of esophageal cancer progression. The main contributions of this work include:

1. DL-based approach: We present a DL-based framework that integrates gene selection with enrichment analysis for examining gene expression data. This approach characterizes the mechanisms underlying esophageal cancer drug response. It identifies critical genes, candidate drugs, therapeutic targets, key pathways, and regulatory factors associated with treatment outcomes.
2. Model training: gene expression datasets associated with esophageal cancer treatment responses were obtained from the Gene Expression Omnibus (GEO) database (accession numbers GSE234304 and GSE273848). After preprocessing, a fully connected deep neural network was trained to identify complex nonlinear patterns in the cell expression profiles. This process implicitly reduced the data’s dimensionality, facilitating the identification of the most relevant genes for predicting drug response.
3. Gene weighting and ranking: Gene selection strategies based on deep learning were employed to prioritize genes according to their contribution to model predictions. Specifically, two approaches were implemented: (1) a base neural network (NN) model followed by support vector regression (SVR) and (2) integrated gradients (IG) applied to the trained network followed by SVR. The genes were ranked based on their estimated importance scores derived from each method. The top-ranked genes identified by these approaches were then analyzed using Enrichr and Metascape for enrichment to characterize the biological pathways and processes associated with drug response and resistance in esophageal cancer.
4. Biological results: The proposed DL-based framework demonstrated superior performance compared with baseline methods. Notably, the IG + SVR approach identified the greatest number of expressed genes in established esophageal cancer cell lines, including TE6, KYSE140, OE19, TE10, TT, and TE11. This indicates greater sensitivity in capturing biologically relevant features. Additionally, the framework identified several candidate drugs, including olaparib, nintedanib, and quercetin.

The rest of the paper is organized as follows: Section 2 describes the datasets and the proposed computational framework. Section 3 presents the experimental methodology and results. In Section 4, the findings are discussed in detail, and Section 5 concludes the paper with a summary of key outcomes and directions for future research.

## 2. Materials and Methods

### 2.1. Gene Expression Profiles

In this study, we utilized the two RNA sequencing datasets, obtained from the Gene Expression Omnibus (GEO) under accession number GSE234304 [17] and GSE273848 [18], respectively. The datasets correspond to esophageal tissue samples from patients with esophageal adenocarcinoma and non-tumor tissues. GSE234304 comprises expression profiling by high-throughput sequencing of human (Homo sapiens) samples and includes a total of 49 esophageal normal and adenocarcinoma tissues, while GSE273848 includes a total of 40 esophageal normal and adenocarcinoma tissues. The datasets consist of bulk cell RNA-seq gene expression profiles, where each sample *x_i_* ∈ *R^n^* represents the expression levels of *n* genes measured from esophageal tissue. Tumor status is appended to patient identifiers – *“*T” for tumor, “N” for non-tumor, and “BO”/“BF” for Barrett’s esophagus, and serves as the primary biological condition for comparison. This design enables downstream computational analyses to characterize transcriptional alterations associated with esophageal adenocarcinoma and to investigate links between microbial dysbiosis and immune infiltration patterns. Table 2 provides an overview of the dataset. The datasets exhibited a moderate class imbalance (GSE234304: 18 tumor vs. 24 non-tumor; GSE273848: 22 tumor vs. 18 non-tumor). While the ratios deviate from perfect balance, they fall within acceptable limits for binary classification without inducing significant bias. Consequently, we opted against aggressive resampling techniques to preserve the natural distribution of biological variance, relying instead on stratified sampling during the train-validation split. The only preprocessing adjustment involved excluding Barrett’s esophagus cases to ensure clear separation between tumor and non-tumor groups for subsequent analyses.

**Table 1.**
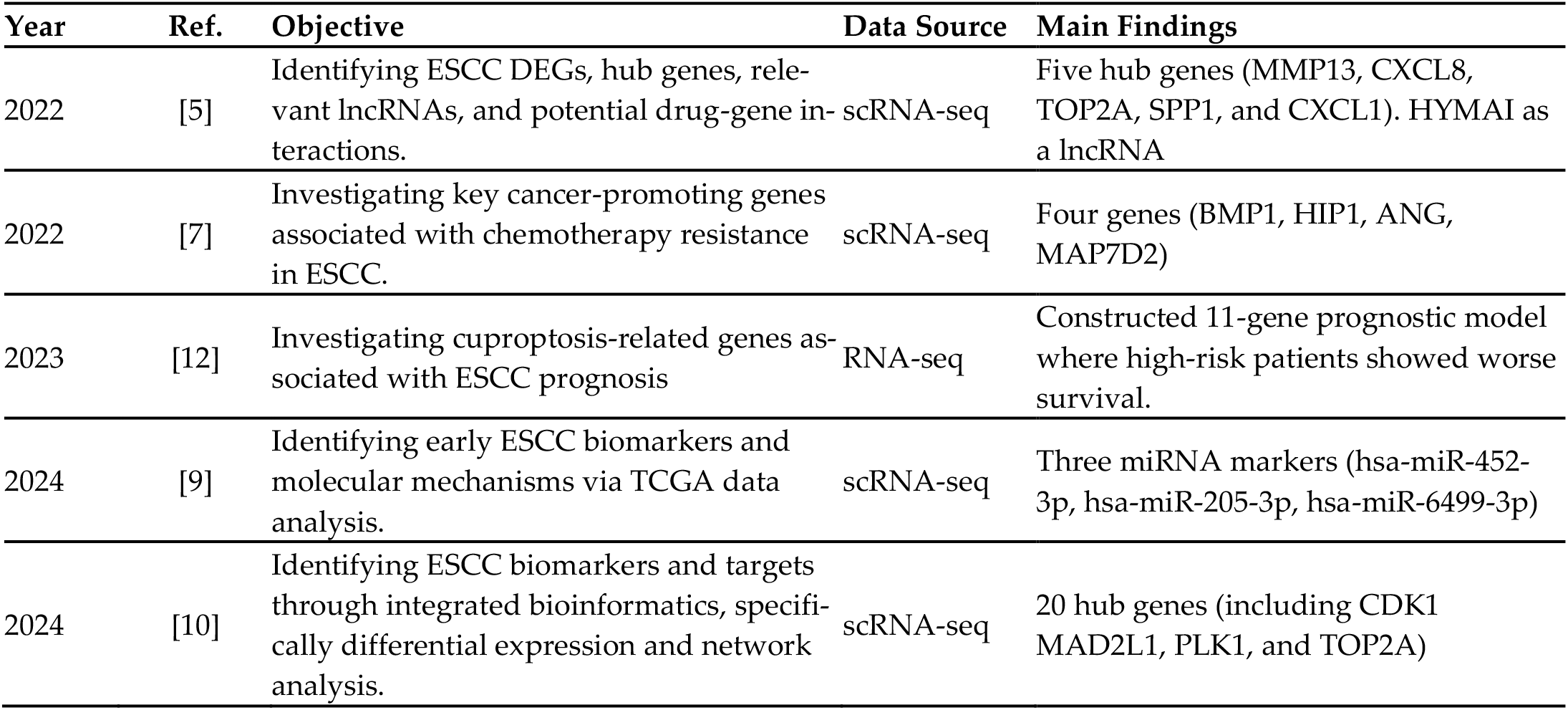
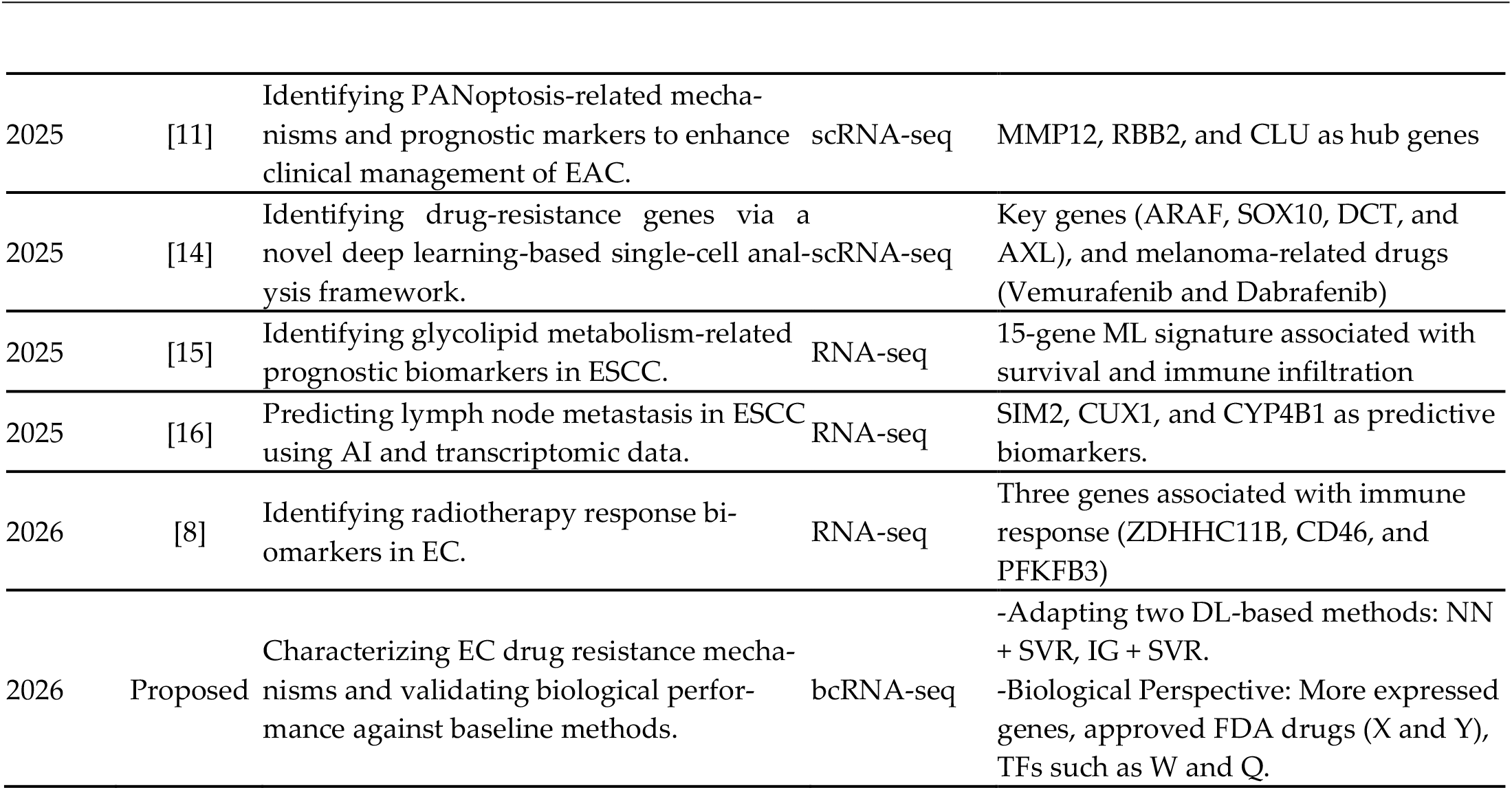
Summary of related work pertaining to Esophageal Cancer compared against our proposed work.

**Table 2.**
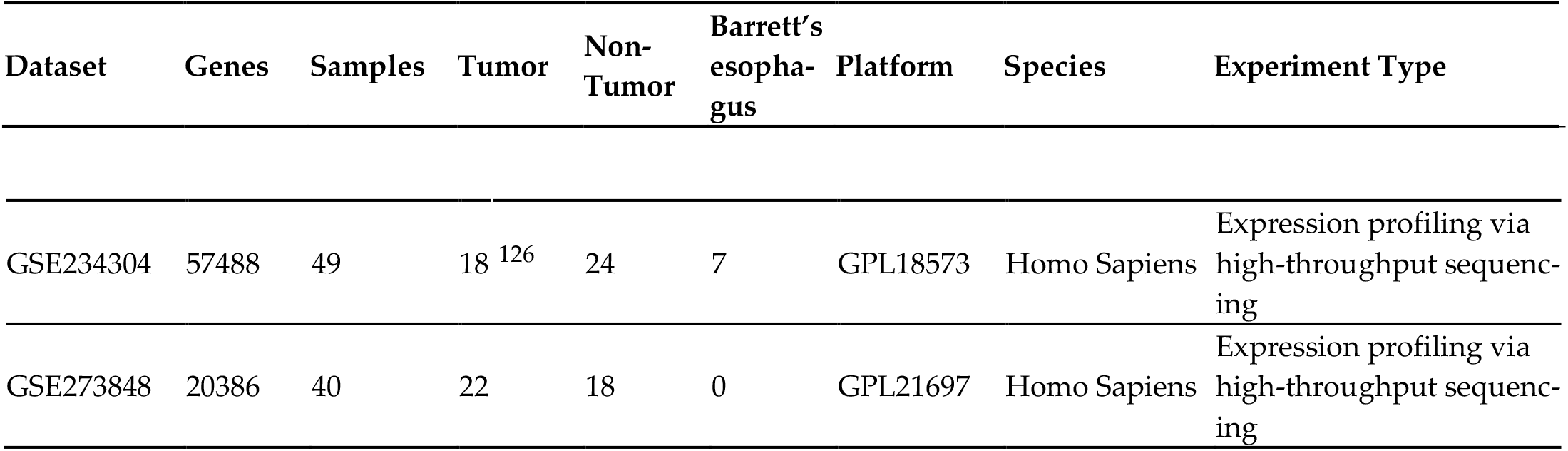
Overview of the bcRNA-seq dataset downloaded from the Gene Expression Om-<u>nibus (GEO) database.</u>

| Dataset | Genes | Samples | Tumor | Non-Tumor | Barrett’s esophagus | Platform | Species | Experiment Type |
| --- | --- | --- | --- | --- | --- | --- | --- | --- |
| GSE234304 | 57488 | 49 | 18 | 24 | 7 | GPL18573 | Homo Sapiens | Expression profiling via high-throughput sequencing |
| GSE273848 | 20386 | 40 | 22 | 18 | 0 | GPL21697 | Homo Sapiens | Expression profiling via high-throughput sequencing |

### 2.2. Computational Approach

The proposed framework encompassing data preprocessing followed by gene ranking using deep learning–based strategies. Specifically, two approaches were implemented; A baseline neural network (NN), after which Support Vector Regression (SVR) was trained on the learned NN weights, and SVR trained on feature attributions derived from the NN using Integrated Gradients (IG). These methods were adapted to bulk gene expression data to identify genes that most strongly contributed to distinguishing tumors from non-tumor samples. The top-ranked genes obtained from each approach were subsequently subjected to enrichment analysis using Enrichr and Metascape to characterize the biological pathways and processes associated with esophageal cancer progression, including mechanisms potentially related to therapeutic response and resistance (Figure 1).

**Figure 1.**
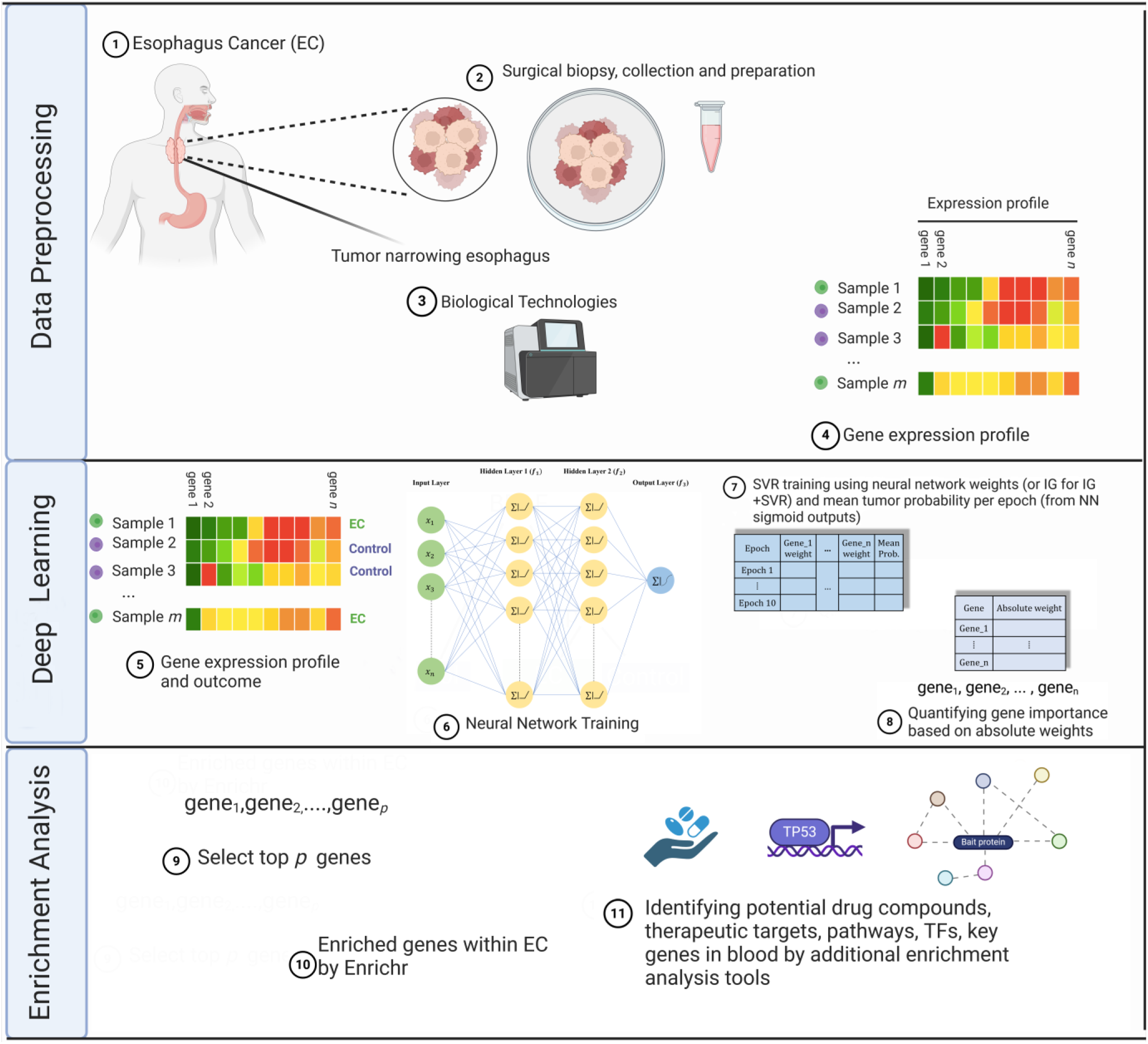
Flowchart of the computational approach identifying drugs, drug targets, critical genes, and transcription factors in esophageal cancer. Data preprocessing: acquiring bcRNA gene expression data corresponding to tumor and non-tumor states in EC from the GEO NCBI database. Deep learning-based gene selection: Computing feature attributions using IG in a neural network provided with bcRNA gene expression data, then used as input features for SVR and subsequently ranking genes from highest to lowest based on the absolute values of the corresponding SVR weights. The top 100 genes identified through this ranking procedure were selected for further analysis. Figure created with BioRender.com. Enrichment analysis: Uploading top 100 genes to Enrichr and Metascape for biological analysis, identifying genes expressed in well-established esophageal cancer cell lines, drugs, drug target, transcription factors, and biological processes and pathways.

#### 2.2.1. Data Preparation

As part of the data preparation process, RNA was extracted from fresh-frozen esophageal tissues, and samples with RNA integrity number (RIN) ≥ 7.0 were selected for sequencing. Sequencing was performed using paired-end reads, followed by alignment to the human reference genome [17].

For the GSE234304 dataset, a total of 42 samples were retained after excluding Barrett’s esophagus cases, which represent precancerous lesions that cannot be definitively classified as either tumor or non-tumor [19], thereby ensuring a clear separation between the comparison groups. One erroneous sample was also excluded from the GSE273848 dataset, leaving a total of 39 samples to be studied. Sample annotations were derived from the pathological status indicated in the patient identifiers, where “N” denoted non-tumor tissue and “T” denoted tumor tissue. Based on this designation, binary labels were assigned to each sample, with 0 representing non-tumor and 1 representing tumor tissues, thereby enabling supervised learning for downstream analyses.

#### 2.2.2. Deep Learning Based Gene Selection

A fully connected feedforward neural network [20] was trained on the bulk cell RNA (bcRNA) sequencing data. Prior to training, min–max scaling was applied to normalize the expression level of each gene to the range [0,1], ensuring comparability across features. The normalized data were provided as input to a fully connected feedforward neural network consisting of two hidden layers with ReLU activation functions [21] and dropout regularization [22] to mitigate overfitting. A two-layer architecture with 128 neurons per layer was chosen. A learning rate of 0.0001 and 10 epochs were determined to be optimal for convergence of the network. The first hidden layer performs a linear transformation defined by a weight matrix *W*_1_ ∈ *R^p^*^×*n*^ and a bias vector *b*_1_ ∈ *R^p^*, where *n* denotes the number of input genes and *p* the number of neurons in the hidden layer (here, *p* = 128). The transformation is followed by a ReLU activation function and a dropout layer with rate 0.1, and the final output layer applies a linear transformation to produce a single logit score, which is subsequently mapped to a probability using the sigmoid function for binary classification (see Figure 2).

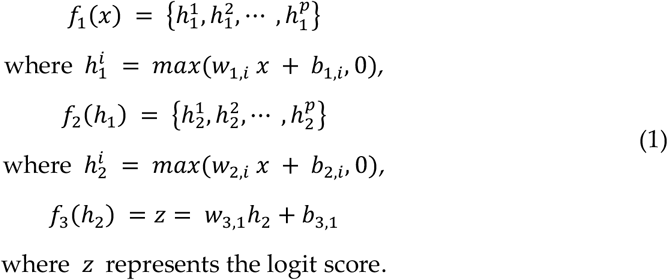

**Figure 2.**
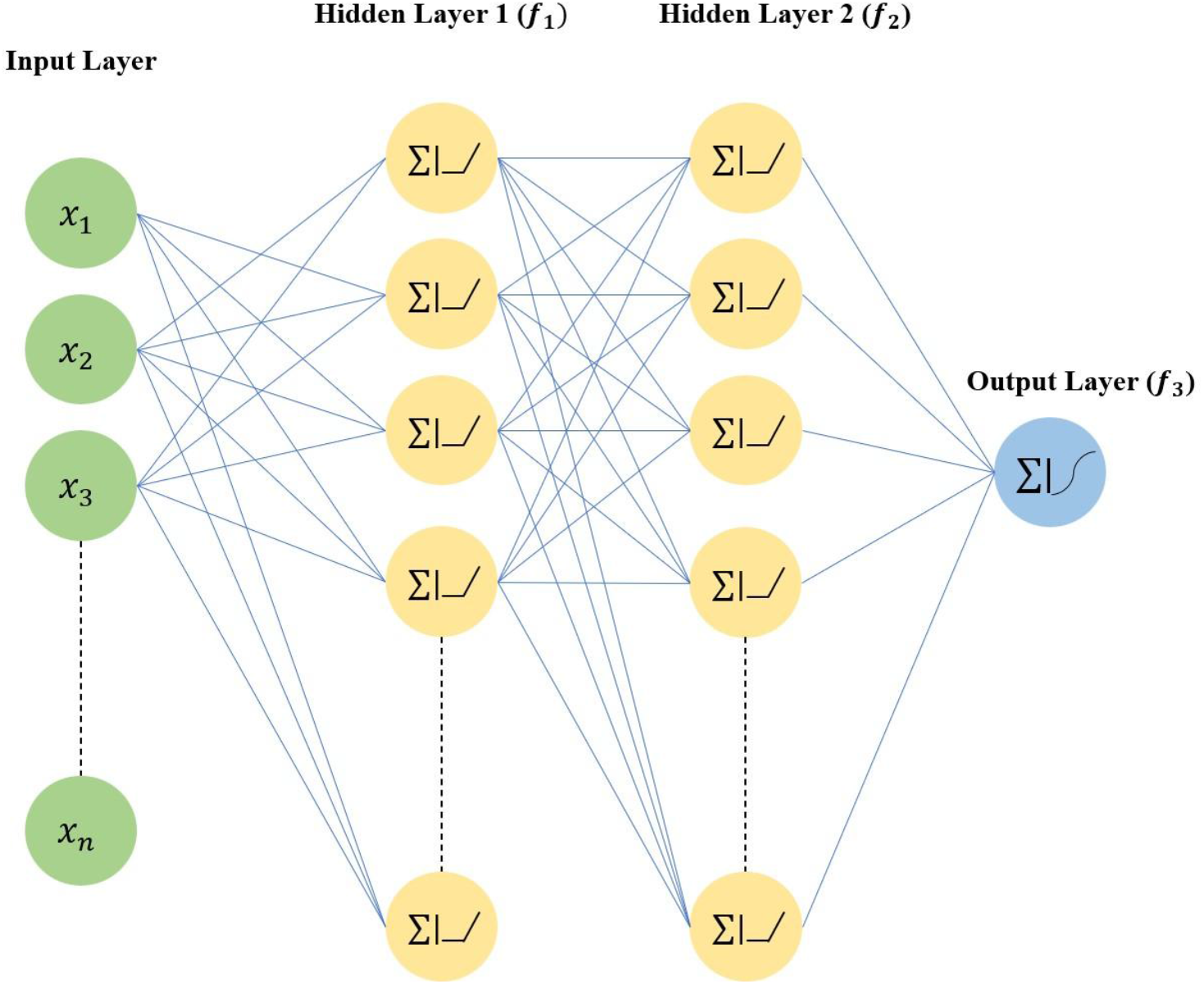
Architecture of the fully connected feedforward neural network in the proposed approach.

where *z* represents the logit score.

During inference, the probability that a given sample belongs to the tumor class is obtained by applying the sigmoid function:

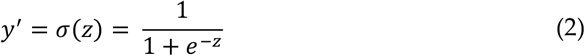

The overall model is thus defined as the composition:

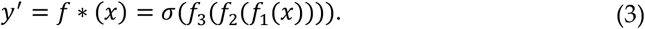

The network was trained using binary cross-entropy with logits loss (BCEWithLogitsLoss), which combines the sigmoid activation and binary cross-entropy into a numerically stable formulation:

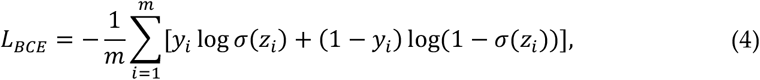

where *m* denotes the number of training samples and *y_i_ ε* {0,1} the true class labels.

Figure 2 visualizing the network architecture, with three weight matrices. The first weight matrix connects the input layer to the first hidden layer, where the gene expression features are used as input. The second matrix links the first hidden layer to the second hidden layer, transforming the intermediate features formed in the first layer. The third weight matrix connects the final hidden layer to the output layer, producing the logit score used for final classification. All network weights and biases were optimized during training using backpropagation in conjunction with gradient-based optimization using the Adam optimizer [23].The resulting weights were stored in an output matrix and used as the input feature set for the SVR.

##### ε-Support Vector Regression (ε-SVR)

To determine the significance of specific genes, we utilized ε-Support Vector Regression (ε-SVR) to model the nonlinear associations within our derived features. Given a training set 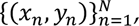 where *x_n_* represents the input feature vector and *y_n_* denotes the observed label, the model is constructed by solving the dual optimization problem. By introducing the dual multipliers *λ_n_* and *λ*^∗^_*n*_, the objective function Q(*λ*, *λ*^∗^) is maximized as shown in Equation 5 [24]:

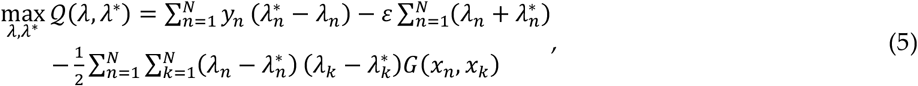

subject to the following constraints:

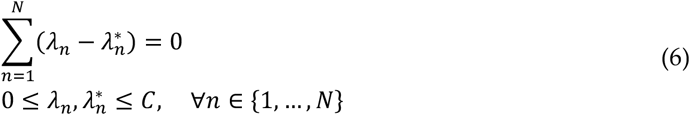

In this formulation, *C* serves as the penalty parameter for deviations exceeding the threshold *ε*. The importance of the input features is interpreted through the weight vector *w* recovered in the transformed feature space:

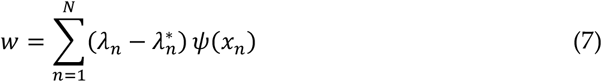

where *ψ*(*x_n_*) represents the high-dimensional mapping induced by the kernel. We specifically employed the Gaussian Radial Basis Function (RBF) kernel, *G*(*x_n_*, *x_k_*), defined as:

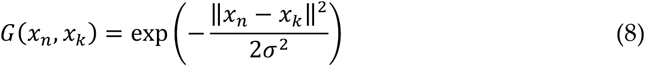

where *σ* modulates the kernel’s reach [25]. The resulting regression surface for any new input *x* is given by:

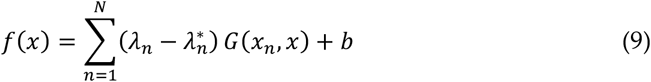

The framework’s data flow is depicted in Figure 3. The input matrix is structured such that rows represent epochs and columns contain the mean gene weights obtained during neural network optimization, alongside a column for the average output logits. Prior to training, all features were standardized to a zero mean and unit variance. The SVR parameters were set to *C* = 10 and *ε* = 0.1, with convergence, reached 50,000 iterations. The model’s predictive reliability was verified via five-fold cross-validation using the *R*^2^ coefficient.

**Figure 3.**
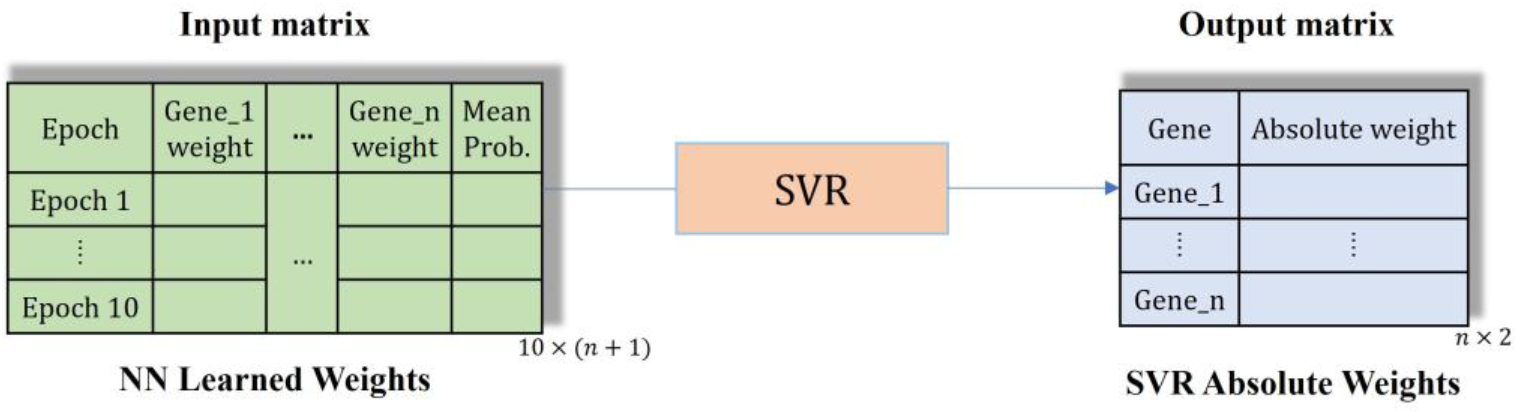
SVR within our pipeline receiving a learned weight matrix and producing an output matrix.

For gene ranking, importance scores were derived from the absolute values of the learned linear coefficients, such that larger absolute weights indicate stronger contributions to the regression function. Genes were then ranked in descending order according to their absolute weights, and the top 100 genes were selected as key contributors to the model’s predictions.

##### Integrated Gradients (IG)

Integrated Gradients (IG) is a gradient-based attribution method [26] used to quantify the contribution of each input feature to a model’s prediction. We applied IG to estimate the relative importance of individual genes in the trained neural network. The method compares the model’s output for a given input to its output for a baseline input, which represents a neutral reference state. The baseline serves as the starting point for computing feature attributions [27]. For a trained model *f*, an input vector *x*, and a baseline input *x*′, the attribution for gene *j* is defined as:

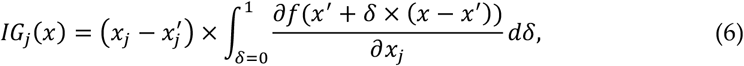

where *δ* ∈ [0,1] interpolates linearly between the baseline *x*′ and the actual input *x*. The integral accumulates the gradients of the model’s output with respect to gene *j* along this path. In our implementation, the baseline was defined as a zero vector, representing the absence of gene expression signal. The integral was approximated numerically using discrete steps along the interpolation path. Rather than directly ranking genes based on raw IG magnitudes, the resulting attribution scores were used as input features for the above-described SVR model. Specifically, for each sample, a vector of absolute attribution values was constructed and standardized. The SVR then learned a linear function:

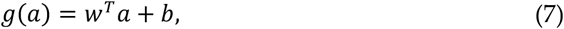

where *a* denotes the attribution feature vector and *w* represents the regression coefficients. Gene importance scores were derived from the absolute values of the learned SVR coefficients:

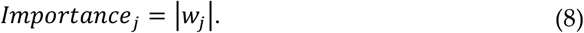

Genes were then ranked in descending order according to these SVR derived weights, and the top 100 genes were selected as key contributors to the model’s predictions.

#### 2.2.3. Enrichment Analysis

To derive biologically meaningful insights, we first trained the aforementioned DL-based methods on the entire dataset, producing 100 genes after ranking them from the highest to the lowest. The most significant genes identified in the cell line were uploaded to Enrichr and Metascape. Gene rankings were determined according to the importance measures produced by each model. The enrichment outputs were then systematically examined to identify biologically relevant terms, including highly expressed genes, candidate drugs, potential therapeutic targets, transcription factors, and related functional categories. The detailed results of this analysis are presented in the following section.

## 3. Results

### 3.1. Experimental Methodology

We compared our DL-based approach with these baseline methods: LIMMA [28], SAM [29], and the t-test [30]. The input provided to each method is gene expression data with binary conditions. In terms of LIMMA, SAM, and t-test baseline methods, statistically significant genes were selected according to adjusted p-values < 0.01. Produced genes via each method were provided to Enrichr (https://maayanlab.cloud/Enrichr/, accessed on 3 February 2026) and Metascape (https://metascape.org/gp/index.html, accessed on 15 February 2026). We retrieved terms related to esophageal cancer in which a method is superior when exhibiting more genes with studied terms. In this study, we utilized Python 3.12.11 to run our DL-based gene selection methods [31]. Specifically, we utilized PyTorch 2.9.1 for neural networks [32]. To identify important genes, we used SciKit library to run IG and SVR [33]. We incorporated dropout regularization, implemented directly within PyTorch, to encourage sparsity in the model’s weights and thereby identifying the most important input genes. All DL-based experiments pertaining to our approach were implemented on Google CoLab with Python using Intel Xeon CPU@2.0 GHz with 32 GB RAM, and Nvidia P100 GPU with 16 GB VRAM. For the bioinformatics tools, we used R 4.5.2 on AMD Ryzen 5@2.4 GHz with 16 GB RAM. Specifically, we employed the LIMMA package in R [28], utilizing the lmFit and eBayes functions [34] for differential expression analysis, and the siggenes package [34] to run SAM. For the t-test, we employed the t-test function within the stats package [30]. To compute adjusted p-values for the LIMMA, SAM, and t-test methods, we employed the p.adjust function with the BH correction.

### 3.2. GSE234304

Table 3 shows the number of expressed genes associated with the retrieved terms (i.e., esophageal cancer cell lines) from Enrichr within the cancer cell line encyclopedia [35]. A higher number of expressed genes indicates superior performance of the computational method. Among all methods used, IG + SVR yielded the highest number of overlapping genes with established esophageal cancer cell lines, with nine expressed genes within the retrieved esophageal cancer cell lines. Specifically, four genes (HSD11B2; GSK3A; DQX1; DEGS2) were expressed within OE19, two genes (PPP6R2; TPCN2) were expressed within TE10, two genes (DQX1; TPCN2) were expressed within TT, and one gene (TPCN2) was expressed within TE11. The second-best method was NN + SVR, obtaining a total of four expressed genes within the retrieved esophageal cancer cell lines; one gene (NENF) was expressed in KYSE150, one gene (SARDH) was expressed in TE8, one gene (ESX1) was expressed in KYSE270, and one gene (ESX1) was expressed in KYSE30. Limma identified three expressed genes within the retrieved esophageal cancer cell lines; one gene (FCF1) within KYSE510, one gene (ANKIB1) within OE33, and one gene (RAB11B) within KYSE30. Neither SAM nor the t-test identified any genes biologically significant to esophageal cancer. In Supplementary Datasheet A1, we include all genes produced via each computational method provided to Enrichr and Metascape. Moreover, Supplementary Table A lists all enrichment analysis results for cancer cell lines encyclopedia obtained from Enrichr.

**Table 3.** Cancer Cell Line Encyclopedia results from Enrichr showing the number of expressed genes associated with each retrieved esophageal cancer cell line, based on gene sets produced by each computational method.

| Method | Rank | Term | Overlap | p-Value | Adjusted p-value |
| --- | --- | --- | --- | --- | --- |
| IG + SVR | 1 | OE19 | 4/272 | 0.047668 | 0.906438 |
|  | 3 | TE10 | 2/108 | 0.101805 | 0.906438 |
|  | 64 | TT | 2/262 | 0.3777 | 0.906438 |
|  | 21 | TE11 | 1/64 | 0.274801 | 0.906438 |
| NN + SVR | 17 | KYSE150 | 1/59 | 0.256339 | 0.892297 |
|  | 48 | TE8 | 1/119 | 0.450234 | 0.892297 |
|  | 51 | KYSE270 | 1/122 | 0.458489 | 0.892297 |
|  | 71 | KYSE30 | 1/161 | 0.555263 | 0.892297 |
| LIMMA | 11 | KYSE510 | 1/46 | 0.206131 | 0.91113 |
|  | 22 | OE33 | 1/71 | 0.299888 | 0.91113 |
|  | 50 | KYSE30 | 1/161 | 0.555263 | 0.91113 |
|  | - | TE11 | 0/64 | - | - |
| SAM | - | OE19 | 0/272 | - | - |
|  | - | TE10 | 0/108 | - | - |
|  | - | TT | 0/262 | - | - |
|  | - | TE11 | 0/64 | - | - |
| t-test | - | OE19 | 0/272 | - | - |
|  | - | TE10 | 0/108 | - | - |
|  | - | TT | 0/262 | - | - |
|  | - | TE11 | 0/64 | - | - |

Figure 4 illustrates the overlap among gene sets identified by each computational method in our framework, alongside those obtained using conventional bioinformatics approaches. A total of 96 distinct genes were identified by each of the NN+SVR and IG+SVR models, whereas all 100 genes reported by LIMMA were unique to it. In contrast, other traditional statistical techniques, including SAM and the t-test, did not identify any significant genes. The degree of overlap between methods was limited, with just four shared genes observed between the NN+SVR and IG+SVR approaches, and no overlap with LIMMA. These findings indicate that the proposed computational models generated distinct gene signatures while identifying a greater number of genes associated with esophageal cancer compared to conventional tools. The complete list of genes corresponding to the UpSet plot in Figure 4 is provided in Supplementary Datasheet A2. The number of overlaps is 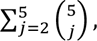 where 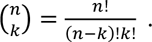

**Figure 4.**
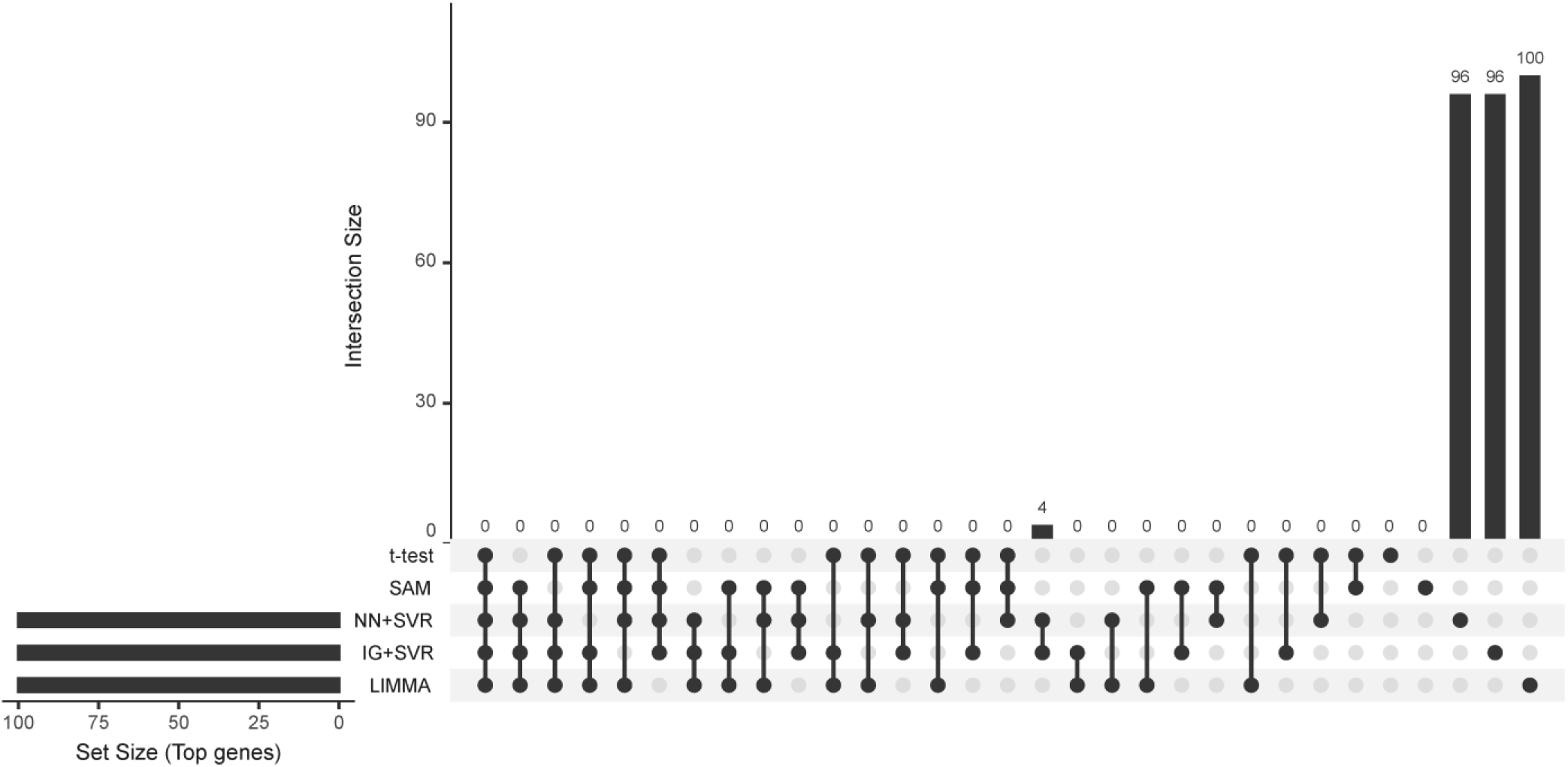
UpSet plot showing overlap among gene lists generated by all computational methods for the esophageal cancer cell line using GSE234304 dataset.

As IG+SVR demonstrated superior performance relative to the other approaches, the top 100 genes identified by this method were further analyzed using Metascape for enrichment analysis. Figure 5a illustrates the protein–protein interaction (PPI) network cluster derived from this analysis. From the constructed PPI network, three key genes were identified: PMPCB, SUMO1, and CEBPB. SUMO1 acts as a key mediator of SUMOylation, regulating transcriptional activity and protein stability in gastrointestinal cancers, including esophageal cancers [36]. SUMO1 has also been identified as a bona fide PLK1-associated target in esophageal cancer and mediates SUMOylation of IκBα to regulate NF-κB signaling and tumor progression in esophageal cancer [37], [38]. CEBPB has been reported to transcriptionally activate MMP3, thereby promoting cancer cell invasion and metastasis in EC [39], [40]. CEBPB also regulates the expression of immunosuppressive genes, including Arg-1, COX2, NOS2, and NOX2, through interaction with Lnc-17Rik, facilitating remodeling of the LAP/LIP/CHOP transcriptional repressor complex and promoting the recruitment of WDR5, leading to increased H3K4me3 enrichment at target gene promoters. As a result, myeloid-derived suppressor cell (MDSC) activity is enhanced, contributing to tumor progression. Notably, the human homolog of CEBPB is significantly overexpressed in MDSCs derived from patients with esophageal cancer [40][41].

**Figure 5.**
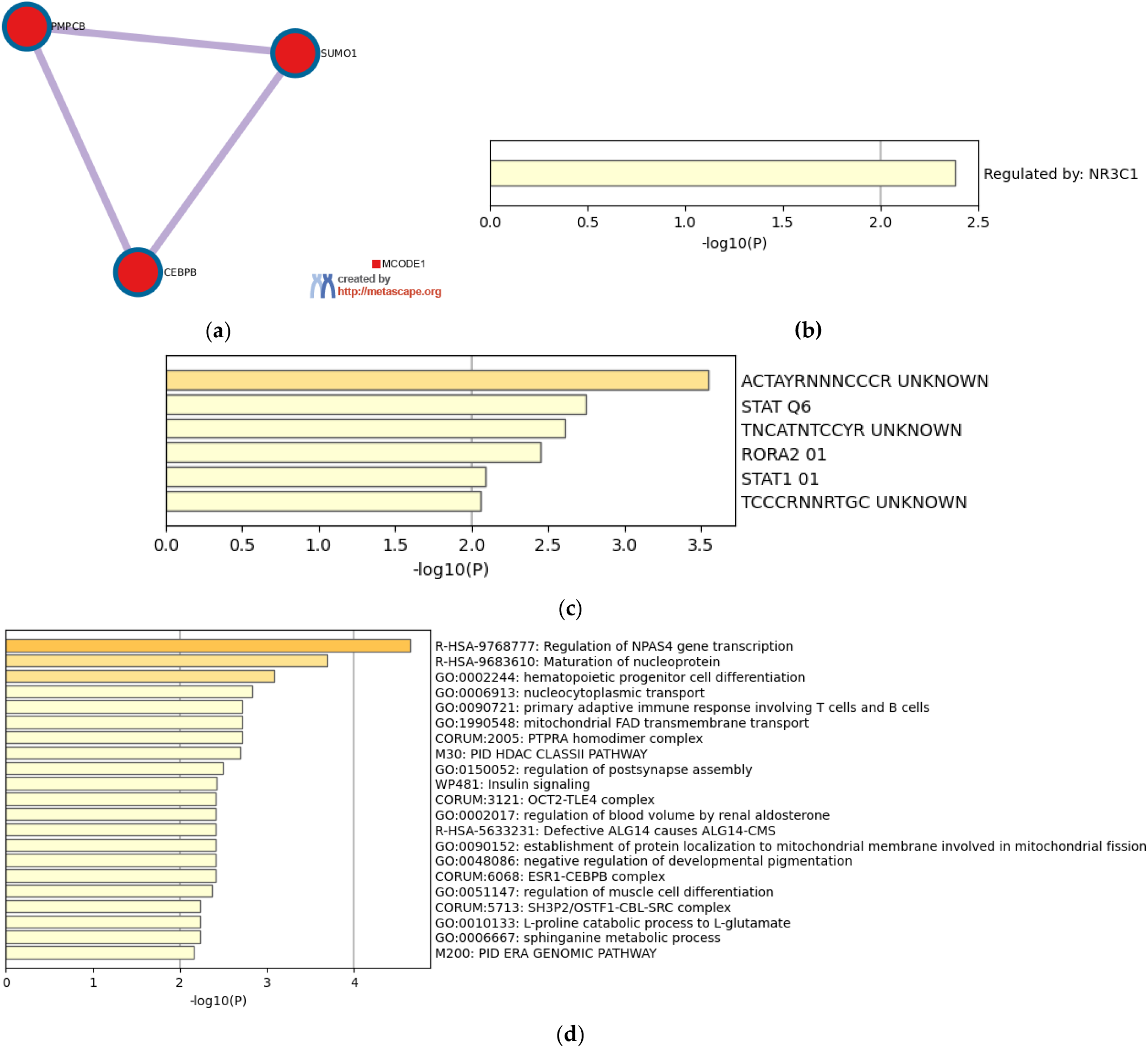
(**a**) A protein–protein interaction (PPI) network cluster identified using Metascape. (**b**) A transcription factor identified through enrichment analysis using Metascape. (**c**) Enrichment results corresponding to transcription factor target genes. (**d**) Biological processes and pathways enriched using Metascape.

Figure 5b presents NR3C1 identified as a transcription factor (TF) using Metascape. NR3C1 encodes the glucocorticoid receptor, and regulates gene expression programs linked to cellular proliferation and inflammatory signaling through interaction with NF-κB and associated co-regulators [42]. In ESCC, NR3C1 has been identified as a predicted target of tumor-derived exosomal miR-103a-2-5p, suggesting its involvement in pathways governing ESCC cell proliferation and migration [43].

Figure 5c displays the transcription factor target enrichment results, highlighting several target gene sets with established relevance to esophageal cancer, including RORA2 and STAT1. The RORA gene family is a circadian clock-associated transcription factor that regulates apoptosis and cell cycle–related pathways in thoracic cancers, including esophageal carcinoma [44]. In EC, RORA functions as a tumor suppressor, where DNMT1-driven promoter methylation reduces its expression under hypoxia, thereby promoting tumor progression through enhanced SLC2A3-mediated glycolysis and metastatic capacity [45]. STAT1 promotes esophageal cancer progression by transcriptionally activating oncogenic targets such as P4HA1, thereby enhancing proliferation and tumor growth [46]. Transcriptomic analyses also identify STAT1 as a core regulatory driver in ESCC shown to repress the tumor suppressor LHPP at the promoter level [47]. Its signaling is closely associated with immune modulation, including correlation with PD-L1 expression and regulation of macrophage polarization, where suppression of STAT1 promotes tumor progression [48], [49].

Figure 5d highlights several enriched biological processes and pathways implicated in esophageal cancer progression. Among the top-ranked terms, nucleoprotein maturation has been associated with disease development. Previous studies have reported enrichment of ribonucleoprotein and ribosomal RNA processing pathways in esophageal cancer. GNL3L, a factor required for pre-rRNA processing during ribosome biogenesis, has been shown to be significantly upregulated in ESCC and correlated with tumor progression and poor clinical outcomes [50], [51], [52]. Hematopoietic progenitor cell differentiation has also been shown to contribute to tumor progression in ESCC, particularly through the expansion and recruitment of myeloid-derived suppressor cells (MDSCs), which arise from immature myeloid progenitors and are significantly increased in ESCC patients, correlating with elevated IL-6 levels, advanced disease stage, and poor prognosis [53], [54], [55]. Insulin signaling is another pathway that is well established to the progression of esophageal cancer. Studies have shown that insulin/IGF receptor activation stimulates the PI3K/AKT/mTOR and RAS/MAPK pathways to promote cell proliferation, survival, and gene transcription [56]. Hyperinsulinemia is also recognized as an oncogenic factor contributing to cancer initiation and progression, as elevated insulin levels promote cellular proliferation and survival while physiological insulin signaling drives broad transcriptional remodeling of metabolic and growth-related gene networks [56], [57]. The complete enrichment analysis results from Metascape have been provided in Supplementary Archive A.

Table 4 summarizes the drugs and their corresponding target genes identified from the IDG Drug Targets 2022 database, with emphasis on their relevance to esophageal cancer treatment and therapeutic resistance. Nintedanib, an orally available triple angiokinase inhibitor targeting VEGFR1–3, FGFR1–3, and PDGFRα/β, suppresses receptor tyrosine kinase signaling pathways that drive angiogenesis and activate downstream PI3K/AKT and MAPK cascades implicated in esophageal cancer progression [58]. In a phase II trial of the same study, nintedanib achieved the predefined 6-month progression-free survival endpoint and demonstrated disease stabilization with manageable toxicity [58]. Moreover, nintedanib is investigated in patients with advanced esophagogastric cancer (https://clinicaltrials.gov/study/NCT02234596) [59]. Quercetin, a naturally occurring flavonoid, inhibits PI3K signaling and modulates downstream survival pathways implicated in esophageal cancer proliferation and therapeutic resistance [60]. It enhances chemotherapy-induced apoptosis through suppression of NF-κB–regulated anti-apoptotic proteins and caspase activation and reverses 5-fluorouracil resistance via modulation of the NRF2/HO-1 axis [61], [62]. Quercetin has also been shown to exert immunomodulatory effects within the tumor microenvironment, supporting its relevance in esophageal cancers characterized by immune and inflammatory pathway enrichment [63].

**Table 4.** Enriched terms from the IDG Drug Targets 2022 dataset identified using Enrichr for the gene set produced by IG+SVR. The table lists associated genes (genes), drug terms (term), and their ranking (rank) in the enrichment results.

| Rank | Term | Class | Genes | Approval Status/Study Phase | References |
| --- | --- | --- | --- | --- | --- |
| 2 | Enoxolone | ERK activator | HSD11B2 | Investigational<br>Preclinical studies | [64] |
| 3 | Verapamil | P-gp inhibitor | TPCN2 | Investigational<br>Preclinical studies | [65], [66] |
| 4 | Quercetin | PI3K-Akt inhibitor | GSK3A | Investigational<br>Preclinical studies | [61], [62], [63] |
| 6 | Nintedanib | VEGFR1–3,<br>FGFR1–3, PDG-FR$\alpha/\beta$ | GSK3A | ClinicalTrials.gov<br>Study | [58], [59] |

### 3.2. GSE273848

The number of expressed genes associated with terms retrieved from the esophageal cancer cell lines have been reported in Table 5. Among all the studied methods, IG + SVR demonstrated superior performance compared to the rest of the methods, identifying a total of 11 expressed genes within well-established esophageal cancer cell lines. Specifically, four genes (BAIAP2L2; CREG2; FAM3B; TNNI3) were expressed within OE19, three genes (CREG2; COX6B2; ABCA13) were expressed within ECGI10, two genes (SOSTDC1; ABCA13) were expressed within KYSE140, and two genes (SOSTDC1; ABCA13) were expressed within TE6. The second-best method was t-test, identifying a total of seven expressed genes within the retrieved esophageal cancer cell lines; three genes (DUOX1; TPRG1; DAPL1) were expressed in KYSE140, three genes (DENND2C; SUSD4; ADH7) were expressed within TE6, and one gene (FAT2) was expressed in ECGI10. SAM identified a total of three expressed genes; one gene (PDPN) was expressed within KYSE140, one gene (PGD) within TE6, and one gene (PLA2G2A) within the OE19 cell line. Limma identified two expressed genes within the retrieved esophageal cancer cell lines; one gene (PACRGL) within KYSE140, and one gene (MAN21C) within TE6. NN + SVR was able to identify only one expressed gene (RNF31) within KYSE140. In Supplementary Datasheet B1, we include all genes produced via each computational method provided to Enrichr and Metascape. Moreover, Supplementary Table B lists all enrichment analysis results for the cancer cell line encyclopedia obtained from Enrichr.

**Table 5.** Cancer Cell Line Encyclopedia results from Enrichr showing the number of expressed genes associated with each retrieved esophageal cancer cell line, based on gene sets produced by each computational method.

| Method | Rank | Term | Overlap | p-Value | Adjusted p-value |
| --- | --- | --- | --- | --- | --- |
| IG + SVR | 25 | ECGI10 | 3/135 | 0.030267 | 0.663788 |
|  | 31 | OE19 | 4/272 | 0.047668 | 0.764260 |
|  | 62 | KYSE140 | 2/102 | 0.092485 | 0.796670 |
|  | 71 | TE6 | 2/110 | 0.104968 | 0.796670 |
| NN + SVR | 192 | ECGI10 | 1/135 | 0.492863 | 0.861934 |
|  | - | OE19 | 0/272 | - | - |
|  | - | KYSE140 | 0/102 | - | - |
|  | - | TE6 | 0/110 | - | - |
| LIMMA | 75 | KYSE140 | 1/102 | 0.401050 | 0.873734 |
|  | 86 | TE6 | 1/110 | 0.424715 | 0.873734 |
|  | - | ECGI10 | 0/135 | - | - |
|  | - | OE19 | 0/272 | - | - |
| SAM | 273 | KYSE140 | 1/102 | 0.401050 | 0.730311 |
|  | 296 | TE6 | 1/110 | 0.424715 | 0.730310 |
|  | 482 | OE19 | 1/272 | 0.746592 | 0.804141 |
|  | - | ECGI10 | 0/135 | - | - |
| t-test | 50 | KYSE140 | 3/102 | 0.014572 | 0.179238 |
|  | 59 | TE6 | 3/110 | 0.017802 | 0.185561 |
|  | 408 | ECGI10 | 1/135 | 0.492863 | 0.741681 |
|  | - | OE19 | 0/272 | - | - |

Figure 6 illustrates the overlap among gene sets identified by each computational method in our framework, alongside those obtained using conventional bioinformatics approaches. A total of 88 distinct genes were identified by each of the NN+SVR and IG+SVR models, whereas 95 of the genes reported by t-test were unique to that method. In contrast, other traditional statistical techniques, including SAM and the LIMMA, identified 97 unique genes. The highest degree of overlap was seen between IG+SVR and NN+SVR with a total of 10 overlapping genes, followed by miniscule overlap amongst other methods. These findings indicate that the proposed IG+SVR computational model generated distinct gene signatures while identifying a greater number of genes associated with esophageal cancer compared to most of the other methods. The complete list of genes corresponding to the UpSet plot in Figure 6 is provided in Supplementary Datasheet B2.

**Figure 6.**
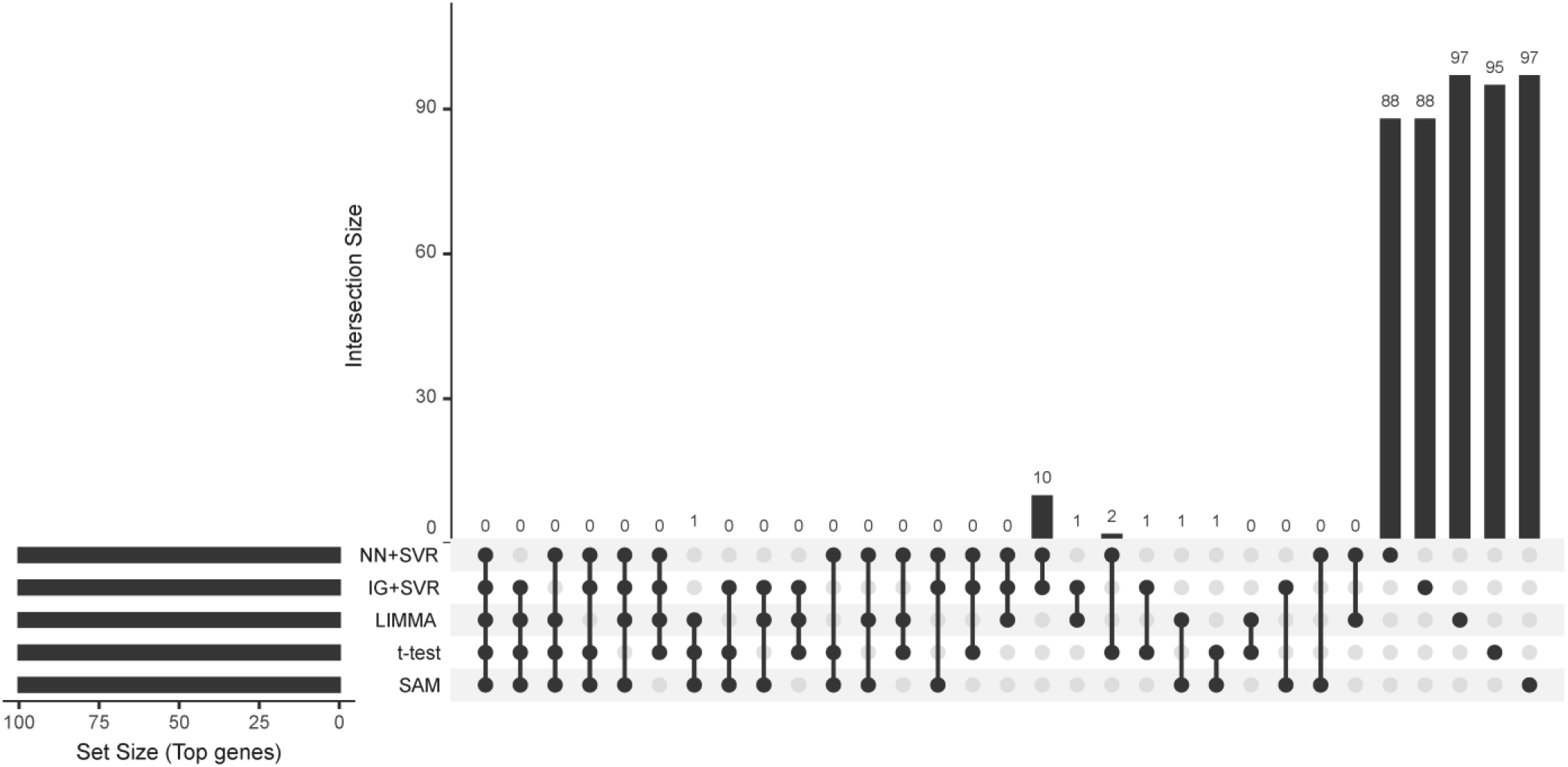
UpSet plot showing the overlap among gene lists generated by all computational methods for the esophageal cancer cell line using GSE273848 dataset.

Since IG+SVR demonstrated superior performance relative to the other approaches and identified genes within a higher number of cell lines, the top 100 genes identified by this method were further analyzed using Metascape for enrichment analysis. Figure 7a presents DDIT3 as a transcription factor identified after Metascape enrichment analysis. DDIT3 encodes CHOP, a canonical pro-apoptotic transcription factor induced by the PERK-eIF2*α*-ATF4 branch of the unfolded protein response during severe ER stress [67]. In esophageal cancer, DDIT3 is consistently identified as a stress-inducible ER-stress/UPR effector that contributes to apoptosis when ER stress is pharmacologically provoked [68], [69]. Another transcription factor identified is RUNX1, which functions as a superenhancer (SE) associated oncogene, promoting proliferation and sitting inside SE-driven transcriptional programs that can be targeted by CDK7 inhibitors [70]. RUNX1 has been shown to often behave as a tumor-suppressive factor in EAC, and to recurrently mutate in familial ESCC [70], [71].

**Figure 7.**
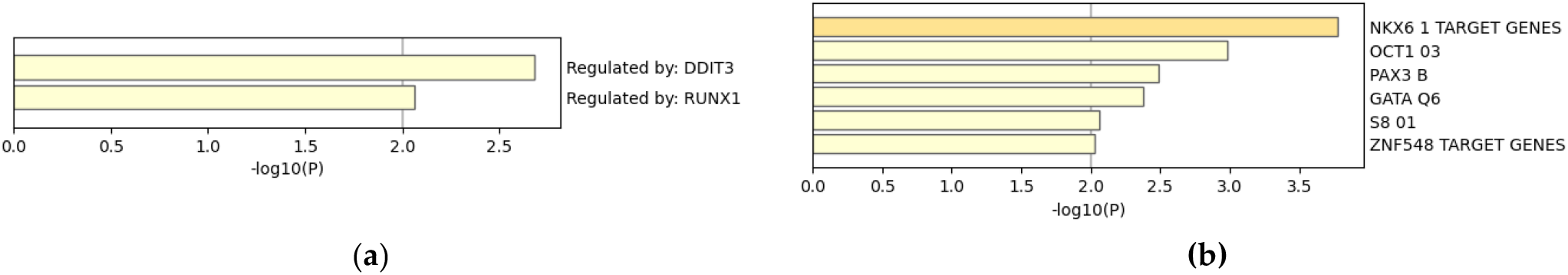
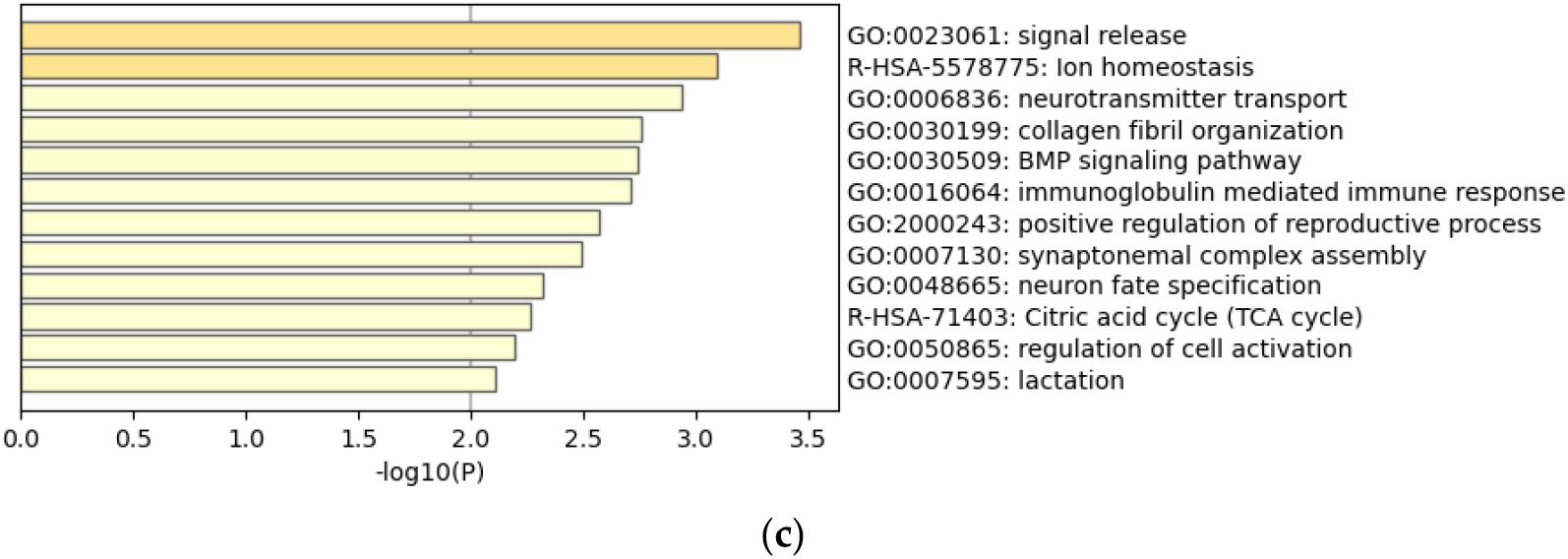
(**a**) Transcription factors identified through enrichment analysis using Metascape. **(b)** Enrichment results corresponding to transcription factor target genes. **(c)** Biological process and pathways enriched using Metascape.

Figure 7b shows the transcription factor targets identified, highlighting several gene sets with relevance to esophageal cancer, including GATA Q6 and OCT1. The GATA Q6 gene has been shown to act as a supervised lineage-survival oncogene in EAC, with amplification/overexpression, survival dependency, and apoptosis signaling via TRAIL/p38*α* [72]. GATA Q6 expression has also been identified to be low or absent in normal squamous epithelium, increased in Barrett’s metaplasia, further increased in dysplasia, and highest in EAC, which is consistent with a role in columnar differentiation and malignant progression [73]. OCT1 overexpression promotes proliferation, colony formation, EMT, migration, and invasion, and its expression is evidently higher in esophageal cancer compared with normal tissue in pan-cancer datasets [74]. Figure 7c highlights a few enriched biological pathways implicated in esophageal cancer progression. Among the top-ranked terms, collagen fibril organization is strongly linked to esophageal cancer progression and prognosis, and gene signatures based on it have been shown to independently predict survival and therapeutic response of patients [75], [76], [77]. BMP signaling has been studied and shown to promote Barrett’s metaplasia, which is an upstream of EAC, and its inhibition shown to reduce Barrett’s epithelium in preclinical models [78], [79]. Studies conducted on the role of TCA in EC tissues show reduced citrate and increased lactate concentrations, with advanced tumors showing further TCA suppression [80]. Metabolic subtyping also shows EAC to be more TCA-dependent than other EC subtypes such as ESCC [81]. Intratumoral immunoglobulin repertoire and *β*-cell infiltration are associated with better EC prognosis and predicted patient response to both chemotherapy and immunotherapy [82], [83]. The complete enrichment analysis results from Metascape have been provided in Supplementary Archive B.

The IDG Drug Targets 2022 database was used to identify drugs corresponding to identified genes, with emphasis on their relevance to esophageal cancer treatment and therapeutic resistance. Table 6 summarizes the identified drugs and corresponding target genes. Olaparib is an oral poly(ADPribose) polymerase (PARP) inhibitor targeting PARP1/2, causing PARP trapping and synthetic lethality in homologous recombination-deficient tumors [84]. In a study, chemotherapy intolerant patients after being switched to olaparib plus nivolumab showed a durable complete response [85]. Olaparib is also being evaluated as a radiosensitizer for EAC patients in a phase II clinical trial [86].

**Table 6.** Enriched terms from the IDG Drug Targets 2022 dataset identified using Enrichr for the gene set produced by IG+SVR. The table lists associated genes (genes), drug terms (term), and their ranking (rank) in the enrichment results.

| Rank | Term | Class | Genes | Approval Status/Study Phase | References |
| --- | --- | --- | --- | --- | --- |
| 15 | Olaparib | PARP1/2 inhibitor | PARP6 | Clinical Trials | [84], [90] |
| 70 | Sunitinib | VEGFR-1/2/3, PDGFR-$\alpha/\beta$ | CAMK2G; MAP4K3 | Clinical Trials | [91], [92] |
| 73 | Pazopanib | VEGFR, PDGFR, c-KIT TKI | MAP4K3 | Pre-clinical Studies | [93] |
| 75 | Sorafenib | VEGFR-2/3, PDGFR-β | MAPK15 | Clinical Trials | [94] |
| 79 | Nintedanib | VEGFR1–3, FGFR1–3, PDGFR$\alpha/\beta$ | MAP4K3 | Clinical Trials | [58], [59] |

Sunitinib is another orally ingested drug, and when combined with irradiation has proven to reduce clonogenic survival, increased apoptosis, increased *γ*-H2AX foci, and altered cell-cycle distribution, thus sensitizing EC cell lines to radiotherapy in vitro [87]. In another study, Sunitinib reduced colony formation and downregulated HIF-1*α* and VEGF, further reducing clonogenic survival, supporting anti-angiogenic and radiosensitizing effects [88]. In an esophageal Barrett’s adenocarcinoma line, Sorafenib has proven to inhibit MAPK-mediated proliferation and reduce ERK phosphorylation and suppressed cell growth [89]. The study also reported common disease stabilization among heavily pre-treated patients who received Sorafenib, indicating some modest activity [89].

## 4. Discussion

To understand the mechanisms underlying drug resistance in esophageal cancer and ultimately improve treatment outcomes and survival, we developed a deep learning (DL)based framework. First, bcRNA-seq gene expression data from esophageal cell lines were retrieved from the GEO repository and used to train a fully connected neural network to distinguish between tumorous and non-tumorous cell lines. The network was optimized using the Adam optimizer with a batch size of 8, a learning rate of 0.0001, and 10 training epochs. Subsequently, a support vector regression (SVR) model was trained on the learned network weights (NN+SVR), and a second SVR model was trained using the network’s integrated gradients (IG+SVR). As described in Section 2.2.2., the top 100 genes with the highest scores were selected for analysis. These top-ranked genes, along with those derived from baseline methods, were subjected to enrichment analysis. Our findings demonstrated that the DL-based approaches identified a greater number of expressed genes across multiple well-established esophageal cancer cell lines compared to conventional methods. Furthermore, these approaches enabled the identification of (1) Clinically studied drugs relevant to esophageal cancer and (2) associated drug targets, transcription factors, biological processes, and signaling pathways.

The results in Section 3 show that IG+SVR, the best-performing method in our framework, identified genes such as SUMO1 and CEBPB, which are known to play critical roles in esophageal cancer progression and immune regulation. SUMO1 has been implicated in regulating transcriptional activity and protein stability through SUMOylation, including modulation of NF-*κ*B signaling and tumor progression in ESCC. Similarly, CEBPB has been associated with enhanced invasion and metastasis through transcriptional activation of oncogenic targets and regulation of immunosuppressive gene expression, contributing to tumor-promoting immune microenvironment remodeling. In addition, transcription factor target analysis identified RORA, GATA, and STAT1, which have established roles in esophageal cancer biology, including regulation of proliferation, malignant progression, inflammatory signaling, metabolic adaptation, and immune modulation. Additionally, our enrichment analysis revealed key biological processes and pathways, including intratumoral immunoglobulin mediated immune response, nucleoprotein maturation, hematopoietic progenitor cell differentiation, and insulin signaling. These pathways are well documented in esophageal cancer research, particularly in the context of tumor progression, ribosome biogenesis, immune cell expansion, and metabolic reprogramming. For example, insulin signaling is crucial for activating PI3K/AKT/mTOR and MAPK cascades, which promote proliferation and survival in esophageal cancer cells. The identification of these genes and pathways highlights the biological relevance and robustness of our DL-based approach.

Our DL-based approach demonstrated clear advantages over traditional bioinformatics tools such as LIMMA, SAM, and the t-test. In both the GSE234304 and GSE273848 datasets, IG + SVR was able to identify more expressed genes in the esophageal cancer cell lines than traditional methods such as LIMMA, SAM and t-test, which even failed to identify any expressed genes in the GSE234304 dataset. These conventional methods rely primarily on linear statistical assumptions and have limited capacity to capture complex, non-linear interactions within high-dimensional transcriptomic data. In contrast, the adapted models within our DL-based framework capture non-linear relationships, enabling the identification of a broader set of biologically relevant genes. Moreover, our analysis highlighted potential drugs and therapeutic candidates relevant to esophageal cancer. Notably, our DL-based approach reduced the search space and identified promising agents such as nintedanib, a triple angiokinase inhibitor targeting VEGFR, FGFR, and PDGFR that suppresses receptor tyrosine kinase signaling implicated in esophageal cancer progression, and quercetin, a flavonoid compound known to inhibit PI3K signaling and modulate NF-*κ*B–mediated survival pathways. While not currently first-line therapies for esophageal cancer, our study suggests they target key pathways that are dysregulated in our identified gene signatures, warranting further investigation for drug repurposing. In addition to these agents, the identified drug-target associations further support the translational relevance of our findings. These results align with emerging therapeutic strategies in esophageal cancer and highlight the potential of our DL-based framework to facilitate drug repurposing efforts. Transcription factor analysis also identified a few genes such as RUNX1 as a tumor-suppressive factor in esophageal cancer, and NR3C1 as a critical regulator of esophageal cancer-associated proliferative and inflammatory signaling programs, underscoring their potential as targets for therapeutic modulation.

Our findings further demonstrate the feasibility of applying deep learning (DL) in the context of drug resistance analysis and highlight the best-performing adapted DL-based feature selection strategy within our framework. Although the training process is computationally intensive, particularly due to the optimization of multiple weight matrices throughout model training, the experimental results support the practicality of DL as a tool for assisting clinicians in prioritizing effective therapeutic candidates during clinical investigation. Notably, identifying important drugs that target key cancer-driving mechanisms in esophageal cancer from a large pool of potential compounds underscores the precision of our approach. Given the vast search space of transcriptomic data and drugtarget databases, identifying clinically relevant candidates is a significant step toward more efficient, targeted therapeutic discovery.

## 5. Conclusions

This study presents a novel deep learning (DL)-based computational framework for identifying critical genes, pathways, therapeutic targets, and drug candidates associated with drug resistance in esophageal cancer. Compared to bioinformatics tools used conventionally, our approach captured a higher number of biologically and clinically relevant genes. Esophageal cancer cell line datasets from the GEO repository were used for validation, demonstrating that the models built using our framework identified a greater number of expressed genes across widely studied esophageal cancer cell lines, such as; OE19, KYSE140, TE6, TE10, TT, and TE11. Our key findings included identification of functionally relevant genes, namely CEBPB, SUMO1, RORA, and STAT1, and transcription factors such as RUNX1 and NR3C1, both of which are implicated in esophageal cancer proliferation, immune modulation, and disease progression. GATA Q6, a transcription factor target shown to have a role in malignant progression of esophageal cancer, was also identified in the study. Additionally, our DL-based framework was able to identify clinically relevant compounds and drugs, such as olaparib, nintedanib and quercetin, highlighting its potential utility in drug repurposing and therapeutic prioritization. These findings underscore the capability of our approach to accelerate the identification of candidate drugs and biologically meaningful targets within large transcriptomic datasets.

Our results demonstrate that the proposed framework can substantially reduce the search space for drugs, therapeutic targets, biomarker genes, biological processes, and pathways associated with esophageal cancer drug resistance. Additionally, it can identify discriminative gene sets with the potential to predict therapeutic response. Our DL-based strategy prioritizes biologically coherent and clinically relevant gene candidates and may serve as a supportive computational tool for clinicians and researchers seeking to streamline therapeutic discovery and precision oncology efforts.

Future work will focus on the following: (1) comparing the performance of different deep and machine learning architectures for classification; (2) applying the framework to identify shared molecular mechanisms between esophageal cancer and other malignancies; (3) determining gene signatures associated with responses to combination targeted therapies; (4) validating these findings in larger, independent cohorts and integrating multi-omics data to improve the framework’s predictive power; and (5) extending the approach to detect significantly mutated genes.

## Supplementary Materials

The following supporting information can be downloaded at: https://www.mdpi.com/article/doi/s1, Datasheet A1: List of top 100 genes produced by each computation method for the GSE234304 dataset; Datasheet A2: Genes corresponding to the UpSet plot for the GSE234304 dataset; Table A: Complete Enrichr Results for the GSE234304 dataset; Archive A: Complete Metascape Results for the GSE234304 dataset; Datasheet B1: List of top 100 genes produced by each computation method for the GSE273848 dataset; Datasheet B2: Genes corresponding to the UpSet plot for the GSE273848 dataset; Table B: Complete Enrichr Results for the GSE273848 dataset; Archive B: Complete Metascape Results for the GSE273848 dataset.

## Author Contributions

F.J.: methodology, software, visualization, investigation, writing—original draft preparation. T.T.: conceptualization, methodology, data curation, supervision, writing—reviewing and editing. F.A.: writing—reviewing and editing. Y.-h.T.: writing—reviewing and editing. All authors have read and agreed to the published version of the manuscript.

## Data Availability Statement

Data is contained within the article.

## Disclaimer/Publisher’s Note

The statements, opinions and data contained in all publications are solely those of the individual author(s) and contributor(s) and not of MDPI and/or the editor(s). MDPI and/or the editor(s) disclaim responsibility for any injury to people or property resulting from any ideas, methods, instructions or products referred to in the content.

